# Vehicle pollution is associated with elevated insect damage to street trees

**DOI:** 10.1101/2022.06.15.496337

**Authors:** Emily K. Meineke, David S. Eng, Richard Karban

## Abstract

1. Vehicle pollution is a pervasive aspect of anthropogenic change across rural and urban habitats. The most common emissions are carbon- or nitrogen-based pollutants that may impact diverse interactions between plants and insect herbivores. However, the effects of vehicle pollution on plant-insect interactions are poorly understood.
2. Here, we combine a city-wide experiment across the Sacramento Metropolitan Area and a laboratory experiment to determine how vehicle emissions affect insect herbivory and leaf nutritional quality.
3. We demonstrate that leaf damage to a native oak species (*Quercus lobata*) commonly planted across the western US is substantially elevated on trees exposed to vehicle emissions. In the laboratory, caterpillars preferred leaves from highway-adjacent trees and performed better on leaves from those same trees.
4. *Synthesis and applications*. Together, our studies demonstrate that the heterogeneity in vehicle emissions across cities may explain highly variable patterns of insect herbivory on street trees. Our results also indicate that trees next to highways are particularly vulnerable to multiple stressors, including insect damage. To combat these effects, urban foresters may consider installing trees that are less susceptible to insect herbivory along heavily traveled roadways.

## Introduction

Trees provide critical ecosystem services to urban residents, such as local cooling, air purification, and runoff reduction (Bolund & Hunhammar, 1999). Most people now live in cities, and global population continues to grow (United Nations, Department of Economic and Social Affairs, Population Division, 2019). Thus, ecosystem service provisioning within urban areas is increasingly important for human health and wellbeing. In addition to the services trees provide to human residents of cities, they also serve as foundational species that support urban biodiversity. In particular, trees are food for insect herbivores with high biomass (caterpillars) and the vertebrates that rely on that biomass, such as birds and lizards.

Herbivores can damage trees by feeding on leaves and reducing photosynthesis (Zangerl et al., 2002) and growth (Zvereva et al., 2012), two functions that are directly correlated with ecosystem services, such as local cooling by trees. Insects can also kill trees (Crawley, 1983), especially when their damage is compounded by abiotic stressors, such as extreme heat and/or drought (Anderegg et al., 2015), which are already prevalent across much of the world’s cities (Tuholske et al., 2021; Zhang et al., 2019). It has long been appreciated that urban trees are under threat from stressors associated with both urbanization and climate change, but the extent to which insect damage may also shift over time is poorly characterized.

The difficulty in predicting the future of insect herbivory in cities largely stems from a poor understanding of the underlying mechanisms driving highly heterogeneous patterns in insect herbivore occurrences and damage within urban areas. Urban habitats are mosaics of heat, nutrients, water availability, and pollution and thus the effects of individual stressors on insect abundance, diversity, and damage are difficult to parse. Given that urban trees are under threat as the climate warms (Burley et al., 2019; Meineke et al., 2016; Meineke et al., 2013; Monteiro et al., 2017), understanding the specific mechanisms driving herbivory damage to urban trees could better prepare planting programs to protect urban trees. An understanding of these mechanisms could also enhance our ability to target tree planting for more optimal insect conservation and management in cities.

Effects of urbanization—and specific aspects of urbanization, such as urban heat—on chewing insects (the most common type of insect damage to plants (Turcotte et al., 2014) are poorly understood. Chewing herbivory increases across some rural to urban gradients (Raupp et al., 2010) and decreases across others. For instance, leaf area damaged by insects was 16.5% higher in rural than in urban areas across several large European cities, an effect attributed to elevated predation by birds and, potentially, ants in city centers (Kozlov et al., 2017). Other studies demonstrate that insect chewing damage is elevated in urban compared to rural areas (Cuevas-Reyes et al., 2013; Dreistadt et al., 1990). Urban to rural gradient studies generally do not reveal the specific mechanisms driving the high heterogeneity of chewing damage within cities.

One underexplored potential mechanism (Leonard & Hochuli, 2017) driving herbivory both within cities and along highways in rural areas is vehicle emissions. The most common vehicle-based pollutants that concentrate within a few hundred meters of roadways are nitrogen oxides (NO_x_), black carbon, carbon monoxide (CO), and ultrafine particulates (Brugge et al., 2007). Many of these compounds are either nitrogen-or carbon-based, and nitrogen and carbon are the backbones of most nutritional and defensive compounds within leaves for insect herbivores. For instance, soluble sugars that serve as nutritional compounds for insects are carbon-based, but so are tannins, which can reduce feeding by herbivores (Barbehenn & Constabel, 2011; Forkner et al., 2004). Similarly, proteins required by insects are nitrogen-based but so are alkaloids, which are compounds that protect against insect herbivory (Macel, 2011; Macel et al., 2005).

Past studies hint at the potential role of vehicle pollution as a driver of leaf nutritional quality for chewing herbivores. At one site in the United Kingdom, trees within 100 meters of motorways were much more likely to be severely defoliated than trees at further distances. Elevated herbivory was attributed to elevated nitrogen dioxide (NO_2_) along highways (Bignal et al., 2007). Similarly, in the Los Angeles Basin, USA, herbivore communities on oak trees at more polluted natural areas tended to be more dominated by chewing herbivores compared to less polluted natural areas (Jones & Paine, 2006). The specific driver of this regional pattern could have been effects of NO_x_ or ozone, and the latter is unlikely to accumulate locally along roadways (Brugge et al., 2007).

Here, we harness the heterogeneity of vehicle pollution across the Sacramento Valley to investigate its effects on insect damage to trees. We employed a spatially explicit model of vehicle emissions (Gately et al., 2015) to select high and low pollution sites across the region, then measured chewing and mining herbivory on trees at these sites to determine relationships between insect damage and exposure to pollutants. Chewing herbivory on leaves is often caused by mandibulate insects with high biomass, notably caterpillars, that serve as food for higher trophic levels and thus could have disproportionate impacts on food webs and ecosystem functions. In the laboratory, we fed naive caterpillars leaves from trees exposed to various levels of pollution to quantify the effects of leaf origin on insect damage to leaves and insect performance. Our results demonstrate that highly polluted, highway-adjacent habitats are associated with shifts in plant-insect interactions and that this topic may be ripe for future research into how roadside environments may affect insect conservation and plant performance in cities.

## Materials and Methods

### Urban field study

#### Description of study region

We examined patterns of herbivory on *Quercus lobata* across the Central Valley of northern California, USA. The region has a Mediterranean climate with monthly mean maximum temperatures ranging between 12.1° C and 33.4° C and monthly mean minimum normal temperatures between 3.6° C and 14.7° C (National Oceanic and Atmospheric Administration, 2022). Roads are not regularly salted, which can affect plant nutritional quality for insect herbivores (Mitchell et al., 2020). However, Sacramento is heavily impacted by vehicle-based pollutants, the most prevalent of which are particulate matter and ground-level ozone (City of Sacramento, 2022).

#### Focal tree selection and sampling

We chose to focus on an oak species because observational studies suggest that interactions between oaks and their herbivores might be affected by pollutants (Bignal et al., 2007; Jones & Paine, 2006). Valley oak (*Quercus lobata*) was ultimately chosen as a focal species because it native, supports a diverse herbivore community of herbivores (Pearse, 2019), and is commonly planted in urban environments across northern California.

Focal trees were selected in QGIS (QGIS Development Team, 2022) by overlaying the 2016 Database of Road Transportation Emissions (DARTE) data layer (Gately et al., 2015) with a shapefile of Sacramento County owned street trees provided by the Sacramento Department of Public Works Urban Forestry Division. The DARTE data layer from 2016 was the latest in a series of one-by-one-kilometer gridded datasets representing vehicle emissions of CO_2_ across the continental United States modelled using traffic data. Though DARTE is meant to represent CO_2_ emissions, the data also represent overall emissions and thus are suitable for describing the emissions of other localized, vehicle-based pollutants.

A five-by-five-kilometer grid was then overlaid on these data layers (Fig. 1a). Within each grid cell, we randomly selected one tree within the highest 10% of polluted zones in the DARTE map, and one tree within the lowest 10% (Fig. 1b). Appropriate trees in each pollution category were not available in all grid cells, but the study design still allowed us to achieve a sample that includes trees in high-(n=14) and low-(n=12) pollution areas across urban Sacramento (Fig. 1c).

**Fig. 1.**
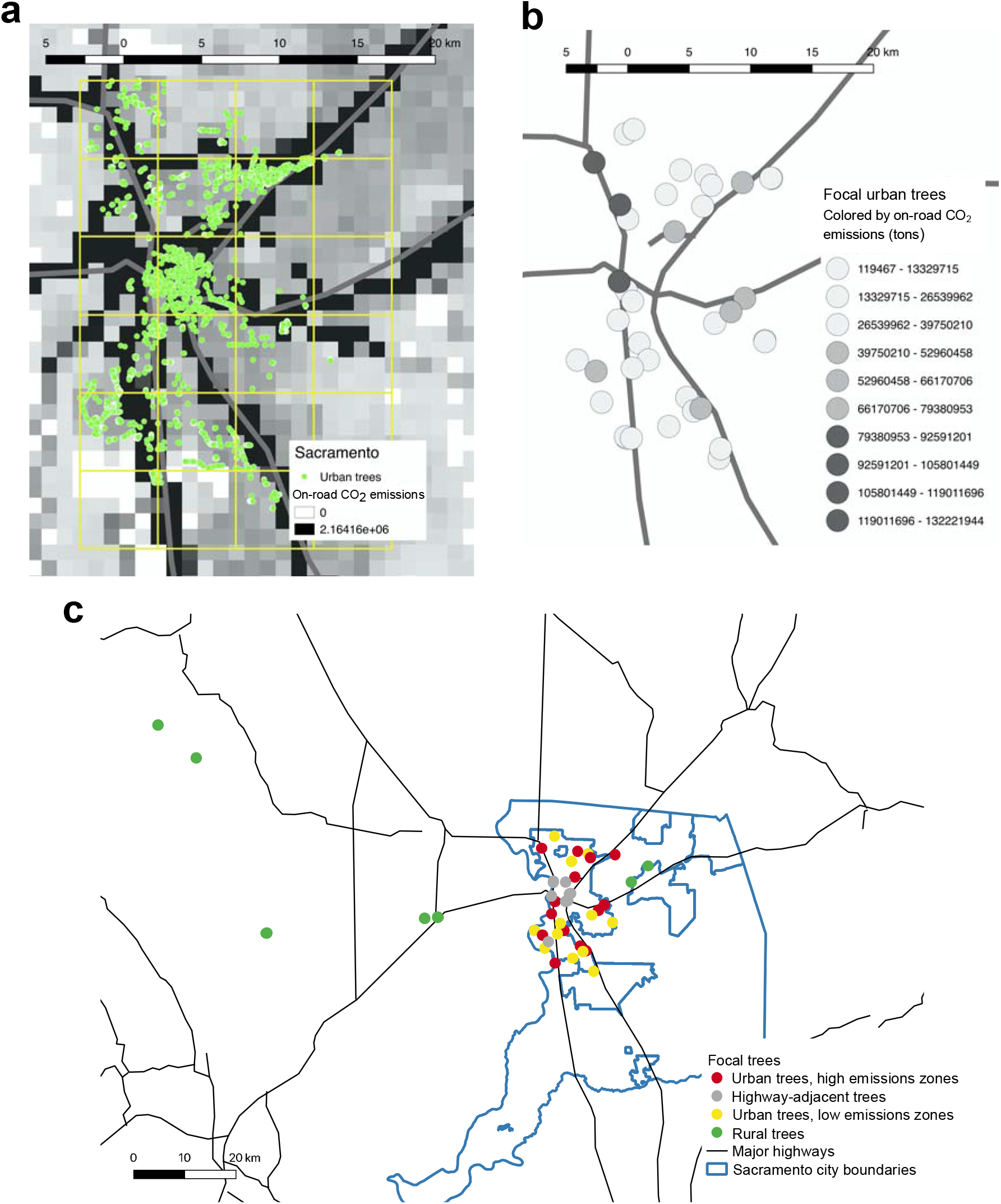
Study system and tree selection in Sacramento, CA. a) The DARTE on-road CO_2_ emissions base layer overlaid with street trees owned by the city of Sacramento City and a 5×5-km grid from which trees in high (top 10%) and low (bottom 10%) vehicle emissions zones were randomly selected. b) Urban focal trees selected from a) and colored by on-road CO_2_ emissions, and c) All focal individuals, including highway-adjacent and rural trees added to study design.

A total of 12 trees from four of the nearest rural sites west of Sacramento were included in the study (Fig. 1c). Three of these trees were from Quail Ridge Reserve and three were from McLaughlin Reserve; these six trees were killed in fires at the end of 2020. In 2021, two additional trees were added to the study from rural regional parks east of Sacramento—which are less likely to be affected by fire—to replace these trees. We added another category of trees we will refer to as highway-adjacent trees, which account for the possibility that resolution of DARTE data (one-by-one kilometer) may not be fine enough to capture the pollution exposure of individual trees because of high levels of air admixture that may dilute pollutants more than a few hundred meters from highways (Brugge et al., 2007). To ensure we included trees exposed to high levels of vehicle pollution in this study, we added 10 focal trees that were accessible from city streets and were within 60 meters of a highway (Fig. 1c). We confirmed with relevant land managers that no focal trees had been treated with pesticides.

One bottom, outer branch was collected from each cardinal direction of each tree with a pole pruner, for a total of four branches per tree. Samples were placed in a plant press, and proportion of leaf area removed by herbivory was estimated by eye on ten leaves per branch in the laboratory. To ensure accurate estimation of herbivory, all participants in the study were trained according to established methods (Johnson et al., 2016). Participants were additionally trained by estimating leaf area removed on a given leaf not included in the study and then using ImageJ (Schneider et al., 2012) to measure leaf area removed. This process was repeated until estimates consistently matched measured values with an error rate of five percent or less. If there were more than ten leaves per branch, focal leaves were randomly selected by overlaying a five-by-five-cm grid on each specimen to avoid any biases for or against damaged leaves. Grid cells were randomly selected, and herbivory was quantified on the topmost leaf within the grid cell. After leaves were selected, the percent leaf area removed by chewing herbivores and the percent leaf area mined by leaf miners was estimated.

#### Covariate measurements

##### Midday water potential

Insect herbivore abundance and herbivory can be affected by plant water stress (Dale & Frank, 2017; Gely et al., 2020; Hahn & Maron, 2018; Huberty & Denno, 2004; Raupp et al., 2010). To estimate water stress of focal trees, we collected three bottom, outer branches from the north side of each tree with a pole pruner between 11:00 and 14:00 between September 15^th^ - 26^th^, 2021. Branches were placed in a pressure chamber (1515D; PMS Instrument Company, Albany, NY, USA), (Dale & Frank, 2017; Meineke et al., 2016).

##### Light pollution

Light pollution can affect herbivory (Mondy et al., 2021) and we measured artificial light at all focal trees. On January 18^th^ - 24^th^, 2022, we use an illuminance meter (T10A; Konica Minolta, Ramsey, NJ, USA) to measure light pollution (Lux) around focal trees. Measurements were taken after the sun had fully set (after approximately 7:30 PM) until 11:00 PM.

##### Diameter at breast height

To account for effects of tree size on herbivory, we measured diameter at breast height at 1.4 meters above the ground in October 2021 after leaf fall.

##### Surface temperature

Because insects are ectothermic, insect physiology, abundances, and damage are often affected by temperature. To account for effects of local temperature on herbivory in this study, we used a surface temperature raster from Landsat 9 Operational Land Imager 2 (OLI-2) (Cook et al., 2014) from August 25, 2022 to extract mean temperatures within a 50-m buffer around each site. Surface temperature layers from other dates and buffer distances (100 meters, 200 meters, 500 meters, and 1000 meters) were included in preliminary models. Those results are not included here because results among the sampled dates and distances were similar.

##### Bulk leaf nutrient

Two leaves per cardinal direction on each tree were collected with a pole pruner, pressed in a plant press, dried and shipped to Brookside Laboratories, INC (New Bremen, Ohio) for analyses of the following nutrients: C, N, P, K, and Na per gm of dry leaf tissue. All nutrients were measured as proportion of leaf weight except for Na, which was measured in parts per million.

##### Normalized differential vegetation index (NDVI)

To account for effects of surrounding vegetation on herbivory, we extracted NDVI from a 200-meter buffer around each tree using an eMODIS NDVI V6 (Earth Resources Observation And Science (EROS) Center, 2002) dataset from May 2020 and calculated average NDVI per pixel. We chose May as a focal month because NDVI within our study region peaks during this month (Turner et al., 2020). Vegetation data layers from other dates and buffer distances (50 meters, 100 meters, 500 meters, and 1000 meters) were included in preliminary models. Results among models with NDVI extracted at other buffer distances were similar to results presented here.

##### DARTE-derived vehicle pollution data

We extracted 2016 on-road emissions data within a 200-meter buffer around each tree and calculated average emissions per pixel. On-road emissions data from other buffer distances (50 meters, 100 meters, 500 meters, and 1000 meters) were included in preliminary models. Results among models with on-road emissions extracted at other buffer distances were similar, and thus are not reported here.

##### Tree proximity to highways

We manually measured the shortest distance from each tree to an Interstate, U.S., or state highway on Google Maps. This variable was included in a subset of models to determine if 1) distance from a major road can serve as a proxy for exposure to vehicle pollution, and 2) on-road emissions explains additional variation not associated with proximity to roads. For instance, if proximity to highways were significant in models, but on-road emissions were not, we could conclude that some other aspect of highway adjacency unrelated to vehicle pollution drives herbivory.

##### Leaf proximity to highways

For highway-adjacent trees, we further characterized leaf exposure to on-road pollution by categorizing each cardinal direction of each tree as “bordering” or “not bordering” the highway.

#### Statistical analyses

We implemented series of Bayesian models in Stan with the *brms* package (Bürkner, 2017) in R (R Core Team, 2019). In all models, mean proportion of leaf area affected by the relevant herbivory type per tree was specified as the response. A beta error structure was specified. Models were each fit with 2000 iterations in four chains, and the initial 1000 iterations were decarded after warm-up. We assumed convergence when Rhat values were equal to one and assessed model fit to the observed data using posterior predictive checks in *brms*. We also calculated the variance explained, i.e., the Bayesian R^2^, for each model using the bayes_R2 function in *brms*. In all models, tree-level covariates included latitude, longitude, DBH, NDVI, water potential, surface temperature, and light pollution.

We generated four sets of models, one of which characterizes herbivory across the categories used to select focal trees (rural, urban low pollution, urban high pollution, highway-adjacent). The second set identified specific mechanisms driving herbivory across these categories. In a third model, we assessed the potential for highly localized effects of pollution on herbivory by testing for the effect of highway adjacency within trees (for details, see below). The fourth set of models focused on leaf nutrients. All continuous predictors were centered on zero to allow comparison between effect sizes within models. For all models that included data from multiple years, we included an interaction effect for year and pollution metric to determine whether the effects of pollution depended on year. We subsequently built separate models for each year. Models were generally structured as described in the formula here with slight deviations as described below.

*Mean proportion of leaf area damaged ∼ Beta(p*_*i*_, *n)*

*f(p*_*i*_) = □ + *β*_*1*_*pollution_metric*_*i*_ × *β*_*2*_*year*_*i*_ + *β*_*3*_*DBH*_*i*_ + *β*_*4*_*latitude*_*i*_

*β*_*5*_*longitude*_*i*_ + *β*_*6*_*NDVI*_*i*_ + *β*_*7*_*light_pollution*_*i*_

*β*_*8*_*surface_temp*_*i*_ + *β*_*9*_*water_potential*_*i*_ + *u*_*I*_

Where *mean proportion of leaf area damaged* is the number of grid cells with chewing damage by mandibulate herbivores *p* on tree *i*. We modelled *p*_*i*_ as a function of the intercept (*a*), *pollution_metric*, the pollution metric *i* depending on the specific model (see below), *year*, the year in which herbivory was sampled (2020 or 2021) on tree *i*, environmental and spatial covariates associated with each tree *i*, and *u*_*i*,_ a grouping factor (random effect) of individual tree characteristics accounting for repeated herbivory sampling in 2020 and 2021.

Model Set 1. *Category-based herbivory models:* We assessed the effects of covariates and sampling category on proportion of leaves affected by chewing and mining by insect herbivores. In these models, ‘pollution _metric’ in the model above is specified as the sampling category of each tree.

Model Set 2. *Mechanistic herbivory models:* We assessed the effects of vehicle pollution and environmental and spatial covariates on insect herbivory. In two models—one with mining and one with chewing herbivory as response variables—we included all covariates listed in the formula above and DARTE-derived vehicle pollution data as the “pollution_metric”. In another set of two models, we assessed the effects of proximity to highways on each type of herbivory. In these models, distance from the closest major highway was specified as the “pollution_metric”, and all covariates were included.

Model Set 3. *Within-tree highway adjacency model*: To determine whether leaves that bordered highways were more or less eaten by insects than leaves on the same trees not bordering highways, we subset the full datasets from each year to include only highway-adjacent trees. In these models, we determined whether the “pollution_metric” of bordering highways (binary: yes/no) affected leaf area removed by mandibulate herbivores in each year. This was the sole fixed effect included in the models. Tree was included in each model as a random effect to account for multiple replicates per tree.

Model Set 4. *Leaf nutrient models:* To assess effects of on-road pollution on leaf nutrients, and C:N ratios (Lincoln et al., 1986; Schäedler et al., 2007) we built separate models for each nutrient following the basic structures of Models 1-2 above. Because C, N, P, and K were measured as proportion of leaf composed of each nutrient, beta error structures were specified for these models. A Gaussian error structure was specified for C:N and the model of Na because it was measured as parts per million. For all leaf nutrient models, the “pollution_metric” specified was DARTE-derived vehicle pollution data as it is the most direct measurement of leaf exposure to pollutants.

### Laboratory studies

To determine effects of pollution on caterpillar preference and performance, we collected several hundred individuals of the locally abundant western tussock moth (*Orgyia vetusta*) from Bodega Marine Reserve on July 16, 2021. While the species is a generalist feeder, the populations we collected from feed on coastal bush lupine (*Lupinus arboreus*), are naïve feeders on *Q. lobata*, and are therefore unlikely to possess adaptations to particular *Q. lobata* phenotypes. During transport, caterpillars were fed *Q. lobata* leaves from local trees on the UC Davis campus. They were then starved for 24 hours before the experiments described below, which were performed on July 19-20, 2021 at ambient temperature and humidity in the lab.

#### Choice assays

To determine the extent to which caterpillars choose to feed on leaves from polluted or less polluted trees, we placed 20 caterpillars in 20 mesh bag arenas with one leaf from each of the trees in the rural, urban low pollution, and highway-adjacent sampling categories. To reduce field effort and ensure feasibility, we chose to eliminate the urban high pollution category. Leaves were labeled by tree with sharpies. After 24 hours, leaves were harvested from the bags and placed in a plant press. We then used ImageJ (Schneider et al., 2012) to quantify proportion of leaf area removed by caterpillars. Any prior damage to leaves was also quantified before the leaves were included in mesh bags, though all leaves chosen for the study were undamaged or minimally damaged by herbivores.

#### No choice assays

To determine caterpillar performance on leaves from polluted and less polluted trees, we placed a leaf from each tree described in the ‘Choice Trials’ in its own individual petri dish. Wet cotton was fastened to the petiole of each leaf to ensure they remained hydrated, and the cotton was re-wetted throughout the experiment. One caterpillar was placed in each petri dish. We repeated this setup on five replicates per focal tree. Caterpillars were weighed at the beginning of the experiment and then after 24 hours of feeding *ad libitum*. Any leaf damage present before caterpillars began to feed was recorded categorically, i.e., as presence of mines, galls, chewing, or scale insects and sooty mold.

#### Statistical analyses

To test our hypotheses, we implemented a series of Bayesian models in Stan with the *brms* package (Bürkner, 2017) in R (R Core Team, 2019). Models were each fit as specified for the field experiment.

#### Choice assays

*Mean proportion of leaf area damaged ∼ overdispersed Beta(p*_*ij*_, *n)*

*f(p*_*ij*_) = □ + *β*_*1*_*tree_category*_*i*_ + *u*_*j*_

To assess effects of pollution on herbivory in choice trials, we modelled *p*_*ij*_ as a function of the intercept (*a*), *tree_category* of tree *i* (rural, urban low emissions, highway adjacent), and arena specified as a random effect (*u*_*j*_). The response variable was specified as total proportion of leaf area removed by caterpillars. We chose a zero-inflated (over-dispersed) beta error structure to account for leaves that remained un-eaten; beta error structures require non-zero values.

#### No choice assays

*Caterpillar weight change ∼ Gaussian(p*_*ij*_, *n)*

*f(p*_*ij*_) = □ + *β*_*1*_*tree_category*_*i*_ + *B*_*2*_*prior_herbivory*_*ij*_ + *u*_*I*_

To assess the effects of pollution on leaf nutritional quality for caterpillars, we modelled *p*_*ij*_ as a function of the intercept (*a*), *tree_category* of tree *i* (rural, urban low emissions, highway adjacent). The response variable was specified as caterpillar weight change from the beginning to the end of the experiment. We included *prior_herbivory* on leaf *j* put in petri dishes as a categorical effect with the following categories: no prior herbivory on leaves, chewing, mining, galls, scale insects/sooty mold. A random effect of tree was included to account for multiple replications per study tree (*u*_*i*_).

## Results

### Urban field study

#### Chewing herbivory

In all models, pollution metrics had a significant positive effect on chewing herbivory. In the categorical model (Model Set 1), highway-adjacent trees displayed more herbivory than trees from any other category (Fig. 2; Table 1a; *ß*= 1.46, *CI*_*95*_ = 0.87– 2.05), with urban trees in low-emissions and high emissions categories displaying intermediate levels of herbivory while rural trees displayed the least, though these values were not statistically distinguishable (Fig. 2; Table 1a). Mechanistic models (Model Set 2) demonstrated positive effects of both on-road vehicle pollution (Fig.3a; Fig. S1a; Table 1b; *ß*= 0.042, *CI*_*95*_ = 0.09– 0.75) and tree proximity to highways (Fig. 3c; Fig. S1c; Table 1c; *ß*= -1.02, *CI*_*95*_ = -2.01– -0.04) on herbivory, such that trees in higher pollution zones and trees closer to highways displayed more chewing damage. Further, leaves from the sides of individual highway-adjacent trees bordering a highway displayed more herbivory than the sides of trees not bordering a highway (Model Set 4; Fig. 4; 2020: *ß*= 0.35, *CI*_*95*_ = 0.04– 0.67; 2021: *ß*= 0.33, *CI*_*95*_ = -0.03– 0.67), though the effect was not as strong in 2021. Chewing herbivory was higher overall in 2020 than in 2021 across all models (Fig. 3; Table 1). In most cases, there were not strong interactive effects between year and pollution metric, indicating that effects of pollutants were similar in 2020 and 2021. However, effects of highway-adjacency in categorical models were higher in 2020 (Table 1a; year × highway-adjacent: *ß*= -0.39, *CI*_*95*_ = -0.73– -0.05), as was the effect of on-road pollution, though this effect was not as pronounced (Fig. S2a-b; year × on-road emissions: *ß*= -0.07, *CI*_*95*_ = -0.20– 0.05). Results from models run separately for each year are included in Tables S1-2. No environmental or spatial covariates explained patterns in chewing herbivory except NDVI and tree size, such that larger trees surrounded by less vegetation displayed more herbivory (Fig. 3a,c; Fig. S1; Table 1).

**Fig. 2.**
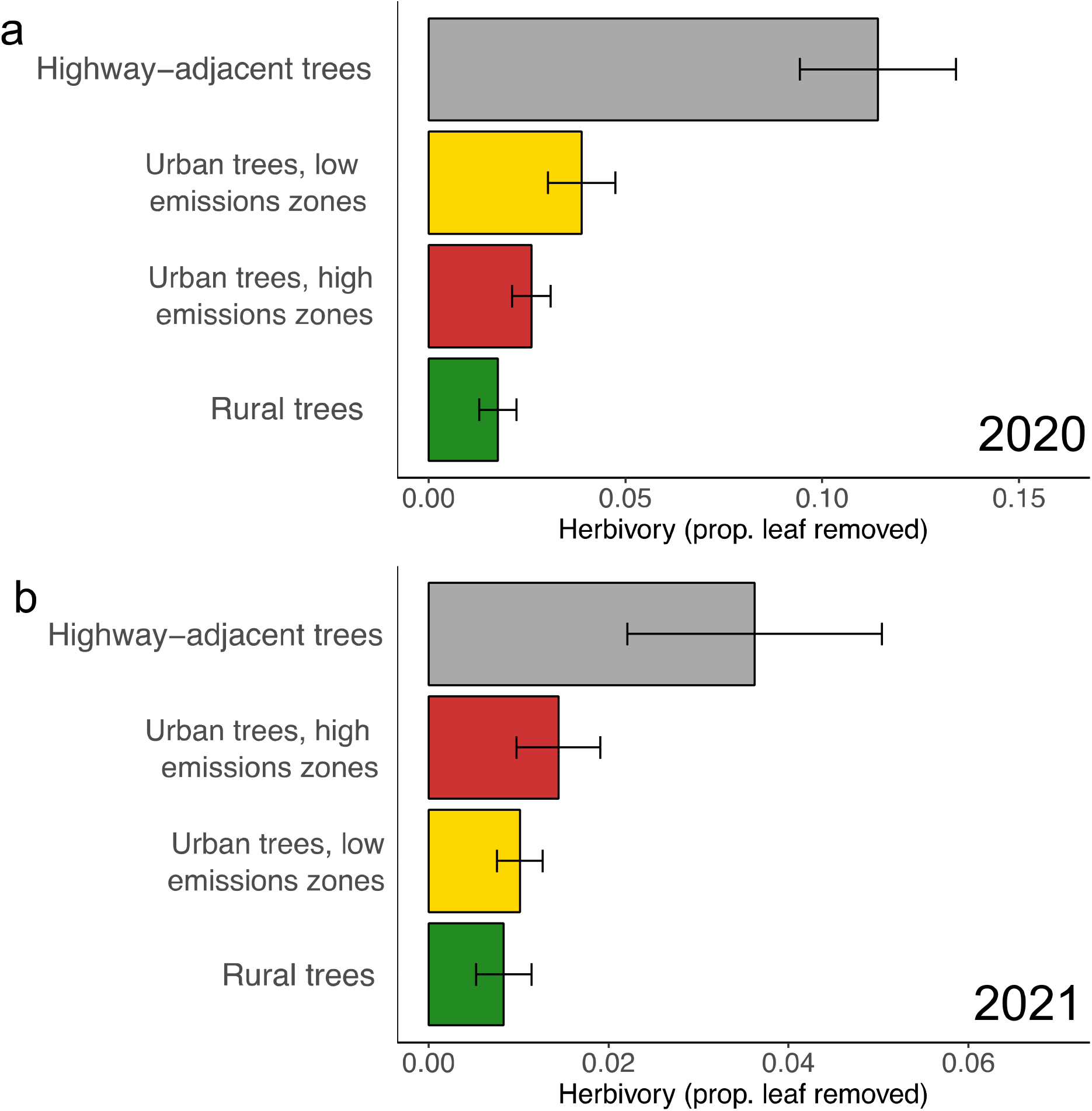
Effects of vehicle-derived pollution on insect herbivory on valley oak (*Quercus lobata*). Chewing herbivory by insect herbivores with mandibles as it relates to categories of pollution exposure.

**Fig. 3.**
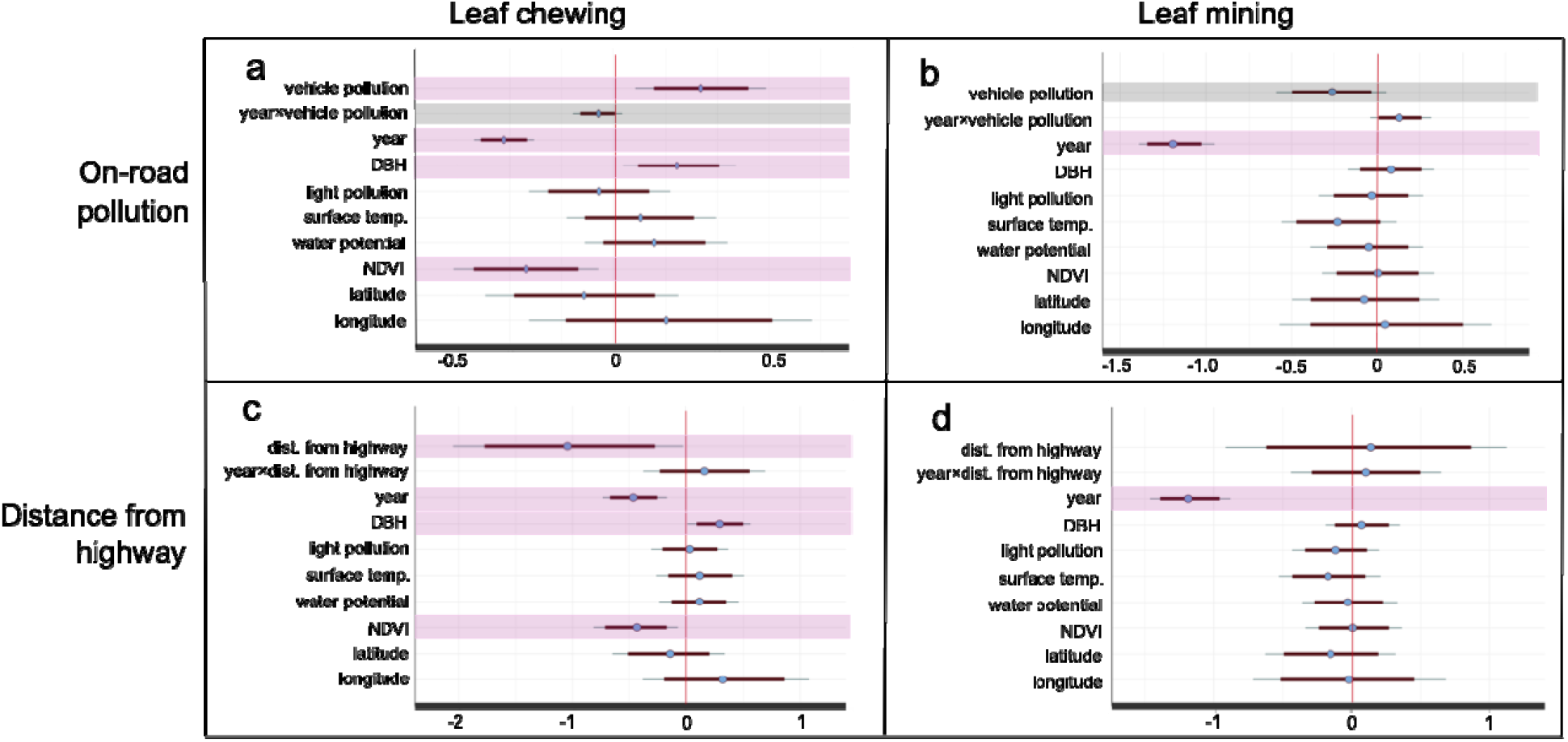
Model estimates showing the predicted effects of pollution, location, and environmental variables on insect damage to leaves. Bold lines represent 85% credibility intervals, and narrow lines represent 95% credibility intervals. Predictors with 85% credibility intervals not crossing zero are highlighted in grey, and those with 95% credibility intervals not crossing zero are highlighted in purple.

**Fig. 4.**
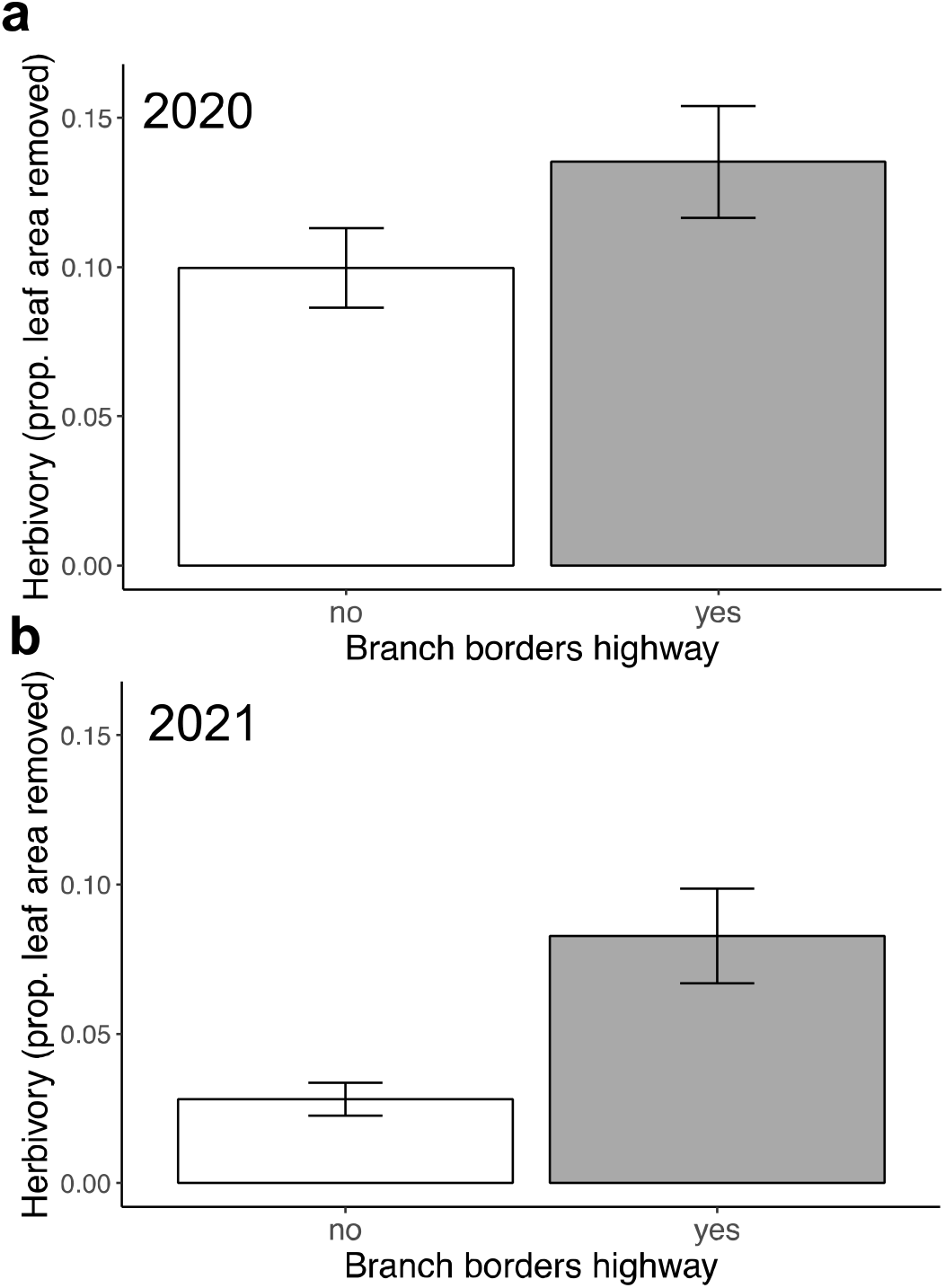
Effects of branch proximity to highways on herbivory. Branches from the sides of individual trees bordering highways displayed higher rates of herbivory than leaves from the same trees not bordering highways.

**Table 1.** Effects of key predictors on insect chewing herbivory. Bayesian models predicting the effects of a) vehicle emissions category, b) distance from the nearest major highway, c) on-road CO_2_ emissions, along with covariates included in all models: light pollution (illuminance), surface temperature, stem water potential (tree water stress), tree size (diameter at breast height), and latitude and longitude of individual focal trees on proportion of leaf area removed by mandibulate herbivores during the 2020 growing season. For each parameter, *ß*_avg_ is the estimated average effect on insect chewing herbivory. Values of each continuous variable were scaled prior to analysis. Thus, larger deviations of *ß*_avg_ values from zero indicate a larger effect of the parameter, and these effects can be compared across parameters. The effective sample size is indicated by n_eff_, and Rhat = 1 indicates convergence.

| Variable | $\beta_{\text{avg}}$ | SE | 2.5% | 97.5% | $n_{\text{eff}}$ | Rhat |
| --- | --- | --- | --- | --- | --- | --- |
| Intercept | -4.11 | 0.30 | -4.71 | -3.50 | 2375 | 1.00 |
| Highway-adjacent | 1.46 | 0.30 | 0.87 | 2.05 | 2200 | 1.00 |
| Urban, low emissions | 0.41 | 0.33 | -0.25 | 1.05 | 2558 | 1.00 |
| Rural | -0.14 | 0.50 | -1.15 | 0.77 | 2684 | 1.00 |
| Illuminance | 0.04 | 0.13 | -0.20 | 0.29 | 3125 | 1.00 |
| Surface temperature | -0.11 | 0.16 | -0.42 | 0.20 | 2834 | 1.00 |
| Water potential | 0.20 | 0.14 | -0.07 | 0.47 | 2415 | 1.00 |
| Tree size | 0.22 | 0.10 | 0.01 | 0.42 | 3172 | 1.00 |
| Latitude | -0.01 | 0.23 | -0.46 | 0.44 | 2467 | 1.00 |
| Longitude | -0.07 | 0.45 | -0.99 | 0.79 | 2365 | 1.00 |
| NDVI | -0.52 | 0.14 | -0.81 | -0.24 | 2941 | 1.00 |
| Year | -0.28 | 0.13 | -0.54 | -0.01 | 2478 | 1.00 |
| Highway-adjacent ×<br>Year | -0.39 | 0.17 | -0.73 | -0.05 | 2894 | 1.00 |
| Urban, low emissions ×<br>Year | -0.39 | 0.20 | -0.78 | 0.01 | 3166 | 1.00 |
| Rural × Year | -0.16 | 0.26 | -0.68 | 0.35 | 3837 | 1.00 |

***b. Mechanistic model including on-road vehicle emissions ( $R^2 = 0.75$ )***
| Variable | $\beta_{\text{avg}}$ | SE | 2.5% | 97.5% | $n_{\text{eff}}$ | Rhat |
| --- | --- | --- | --- | --- | --- | --- |
| Intercept | -3.83 | 0.22 | -4.29 | -3.39 | 1862 | 1.00 |
| On-road CO<sub>2</sub> emissions | 0.42 | 0.17 | 0.09 | 0.75 | 1726 | 1.00 |
| Illuminance | -0.07 | 0.18 | -0.45 | 0.29 | 1868 | 1.00 |
| Surface temperature | 0.12 | 0.18 | -0.24 | 0.46 | 2261 | 1.00 |
| Water potential | 0.19 | 0.18 | -0.15 | 0.54 | 1943 | 1.00 |
| Tree size | 0.31 | 0.14 | 0.04 | 0.59 | 1939 | 1.00 |
| Latitude | -0.15 | 0.24 | -0.61 | 0.32 | 1708 | 1.00 |
| Longitude | 0.28 | 0.36 | -0.41 | 1.02 | 1791 | 1.00 |
| NDVI | -0.41 | 0.18 | -0.76 | -0.07 | 2124 | 1.00 |
| Year | -0.53 | 0.08 | -0.67 | -0.38 | 3076 | 1.00 |
| On-road CO<sub>2</sub> emissions<br>× Year | -0.07 | 0.06 | -0.20 | 0.05 | 4316 | 1.00 |

***c. Mechanistic model including distance from nearest major highway ( $R^2 = 0.73$ )***
| Variable | $\beta_{\text{avg}}$ | SE | 2.5% | 97.5% | $n_{\text{eff}}$ | Rhat |
| --- | --- | --- | --- | --- | --- | --- |
| Intercept | -4.19 | 0.29 | -4.77 | -3.64 | 2033 | 1.00 |
| Distance from highway | -1.02 | 0.50 | -2.01 | -0.04 | 1968 | 1.00 |
| Illuminance | 0.03 | 0.17 | -0.29 | 0.37 | 1892 | 1.00 |
| Surface temperature | 0.12 | 0.19 | -0.25 | 0.48 | 2129 | 1.00 |
| Water potential | 0.11 | 0.18 | -0.23 | 0.46 | 1889 | 1.00 |
| Tree size | 0.30 | 0.14 | 0.02 | 0.60 | 1902 | 1.00 |
| Latitude | -0.15 | 0.25 | -0.65 | 0.34 | 1893 | 1.00 |
| Longitude | 0.33 | 0.37 | -0.38 | 1.07 | 2008 | 1.00 |
| NDVI | -0.44 | 0.19 | -0.81 | 0.08 | 1958 | 1.00 |
| Year | -0.46 | 0.14 | -0.74 | -0.17 | 2198 | 1.00 |
| Distance from highway ×<br>Year | 0.16 | 0.27 | -0.37 | 0.69 | 2473 | 1.00 |

#### Mining herbivory

Mining and chewing herbivory were of similar magnitudes with an average across all trees of four percent of leaf area mined and five percent of leaf area chewed in 2020, and 0.3 percent of leaf area mined and two percent of leaf area chewed in 2021. Models of mining herbivory did not display strong effects of vehicle pollution. However, trends pointed to potential negative effects of pollution on leaf mining. While neither sampling category (Table 2a) nor distance from highways affected mining herbivory (Table 2c; Fig. S3c-d), on-road emissions had a marginally negative effect on mining (Table 2b; Fig. S3a-b; *ß*= -0.26, *CI*_*95*_ = - 0.59– 0.08). Results from models run separately for each year are included in Table S3-4. No additional covariates explained patterns in mining herbivory (Fig. S3; Table 2).

**Table 2.** Effects of key predictors on insect leaf mining herbivory. Bayesian models predicting the effects of a) vehicle emissions category, b) distance from the nearest major highway, c) on-road CO_2_ emissions, along with covariates included in all models: light pollution (illuminance), surface temperature, stem water potential (tree water stress), tree size (diameter at breast height), and latitude and longitude of individual focal trees on proportion of leaf area removed by mining herbivores during the 2020 growing season. For each parameter, *ß*_avg_ is the estimated average effect on insect mining herbivory. Values of each continuous variable were scaled prior to analysis. Thus, larger deviations of *ß*_avg_ values from zero indicate a larger effect of the parameter, and these effects can be compared across parameters. The effective sample size is indicated by n_eff_, and Rhat = 1 indicates convergence.

| Variable | $\beta_{\text{avg}}$ | SE | 2.5% | 97.5% | $n_{\text{eff}}$ | Rhat |
| --- | --- | --- | --- | --- | --- | --- |
| Intercept | -4.43 | 0.41 | -5.24 | -3.64 | 2407 | 1.00 |
| Highway-adjacent | -0.32 | 0.41 | -1.14 | 0.47 | 2638 | 1.00 |
| Urban, low emissions | 0.17 | 0.41 | -0.67 | 0.98 | 2325 | 1.00 |
| Rural | -0.21 | 0.65 | -1.57 | 0.99 | 2713 | 1.00 |
| Illuminance | -0.07 | 0.15 | -0.36 | 0.22 | 2744 | 1.00 |
| Surface temperature | -0.22 | 0.19 | -0.58 | 0.15 | 2591 | 1.00 |
| Water potential | 0.06 | 0.18 | -0.30 | 0.39 | 2354 | 1.00 |
| Tree size | 0.09 | 0.13 | -0.17 | 0.35 | 3012 | 1.00 |
| Latitude | 0.03 | 0.27 | -0.49 | 0.55 | 2192 | 1.00 |
| Longitude | -0.26 | 0.59 | -1.49 | 0.87 | 2609 | 1.00 |
| NDVI | 0.02 | 0.17 | -0.31 | 0.38 | 2787 | 1.00 |
| Year | -1.20 | 0.15 | -1.51 | -0.90 | 2145 | 1.00 |
| Highway-adjacent × Year | 0.34 | 0.23 | -0.10 | 0.78 | 3676 | 1.00 |
| Urban, low emissions × Year | -0.21 | 0.20 | -0.61 | 0.18 | 3911 | 1.00 |
| Rural × Year | -0.07 | 0.22 | -0.52 | 0.36 | 3663 | 1.00 |

***b. Mechanistic model including on-road vehicle emissions ( $R^2 = 0.83$ )***
| Variable | $\beta_{\text{avg}}$ | SE | 2.5% | 97.5% | $n_{\text{eff}}$ | Rhat |
| --- | --- | --- | --- | --- | --- | --- |
| Intercept | -4.57 | 0.23 | -5.01 | -4.10 | 1105 | 1.00 |
| On-road CO<sub>2</sub> emissions | -0.26 | 0.17 | -0.59 | 0.08 | 2557 | 1.00 |
| Illuminance | -0.04 | 0.15 | -0.34 | 0.26 | 2298 | 1.00 |
| Surface temperature | -0.23 | 0.17 | -0.56 | 0.09 | 2310 | 1.00 |
| Water potential | -0.05 | 0.16 | -0.37 | 0.26 | 2143 | 1.00 |
| Tree size | 0.08 | 0.12 | -0.17 | 0.32 | 2321 | 1.00 |
| Latitude | -0.08 | 0.22 | -0.50 | 0.36 | 2104 | 1.00 |
| Longitude | 0.04 | 0.30 | -0.55 | 0.62 | 1975 | 1.00 |
| NDVI | 0.01 | 0.17 | -0.31 | 0.33 | 2270 | 1.00 |
| Year | -1.19 | 0.12 | -1.39 | -0.91 | 711 | 1.00 |
| On-road CO<sub>2</sub> emissions<br>× Year | 0.13 | 0.09 | -0.04 | 0.32 | 3553 | 1.00 |

***c. Mechanistic model including distance from nearest major highway ( $R^2 = 0.83$ )***
| Variable | $\beta_{\text{avg}}$ | SE | 2.5% | 97.5% | $n_{\text{eff}}$ | Rhat |
| --- | --- | --- | --- | --- | --- | --- |
| Intercept | -4.52 | 0.29 | -5.07 | -3.96 | 1474 | 1.00 |
| Distance from highway | 0.14 | 0.52 | -0.88 | 1.15 | 2053 | 1.00 |
| Illuminance | -0.11 | 0.17 | -0.44 | 0.22 | 1935 | 1.00 |
| Surface temperature | -0.17 | 0.19 | -0.54 | 0.21 | 1922 | 1.00 |
| Water potential | -0.02 | 0.17 | -0.37 | 0.31 | 1813 | 1.00 |
| Tree size | 0.08 | 0.14 | -0.20 | 0.35 | 2063 | 1.00 |
| Latitude | -0.15 | 0.24 | -0.61 | 0.31 | 2086 | 1.00 |
| Longitude | -0.03 | 0.34 | -0.71 | 0.64 | 1716 | 1.00 |
| NDVI | 0.01 | 0.18 | -0.33 | 0.37 | 1854 | 1.00 |
| Year | -1.18 | 0.15 | -1.47 | -0.87 | 1561 | 1.00 |
| Distance from highway ×<br>Year | 0.10 | 0.28 | -0.48 | 0.65 | 3090 | 1.00 |

#### Leaf nutrients

We found no evidence for effects of pollution metrics on leaf nutrients (Model Set 4; Fig. S4). The sole covariate with effects on leaf nutrient concentrations was surface temperature, such that trees at sites with higher surface temperatures had marginally lower nitrogen content (Fig. S4d; Table S5a; *ß*= -0.08, *CI*_*95*_ = -0.20– 0.03), and higher C:N ratios (Fig. S4f; Table S5b; *ß*= 2.07, *CI*_*95*_ = 0.03– 4.07).

### Laboratory studies

In choice assays, highway adjacent trees were eaten more than trees from rural areas (Fig. 5a; *ß*= -0.39, *CI*_*95*_ = -0.64– -0.14) and low emissions zones within cities (Fig. 5a; *ß*= -0.44, *CI*_*95*_ = -0.65–0.22). In no-choice assays, most caterpillars lost weight from the beginning to the end of the experiment, but individuals fed leaves from highway-adjacent trees lost less weight than those from rural areas (Fig. 5b; *ß*= -0.01, *CI*_*95*_ = -0.03– 0.00) and low emissions zones within cities (Fig. 5b; *ß*= -0.01, *CI*_*95*_ = -0.03– 0.01), though the latter trend had less support than the former as the credible interval overlapped zero. There was no effect of previous leaf herbivory on caterpillar weight change during the course of the assay.

**Fig. 5.**
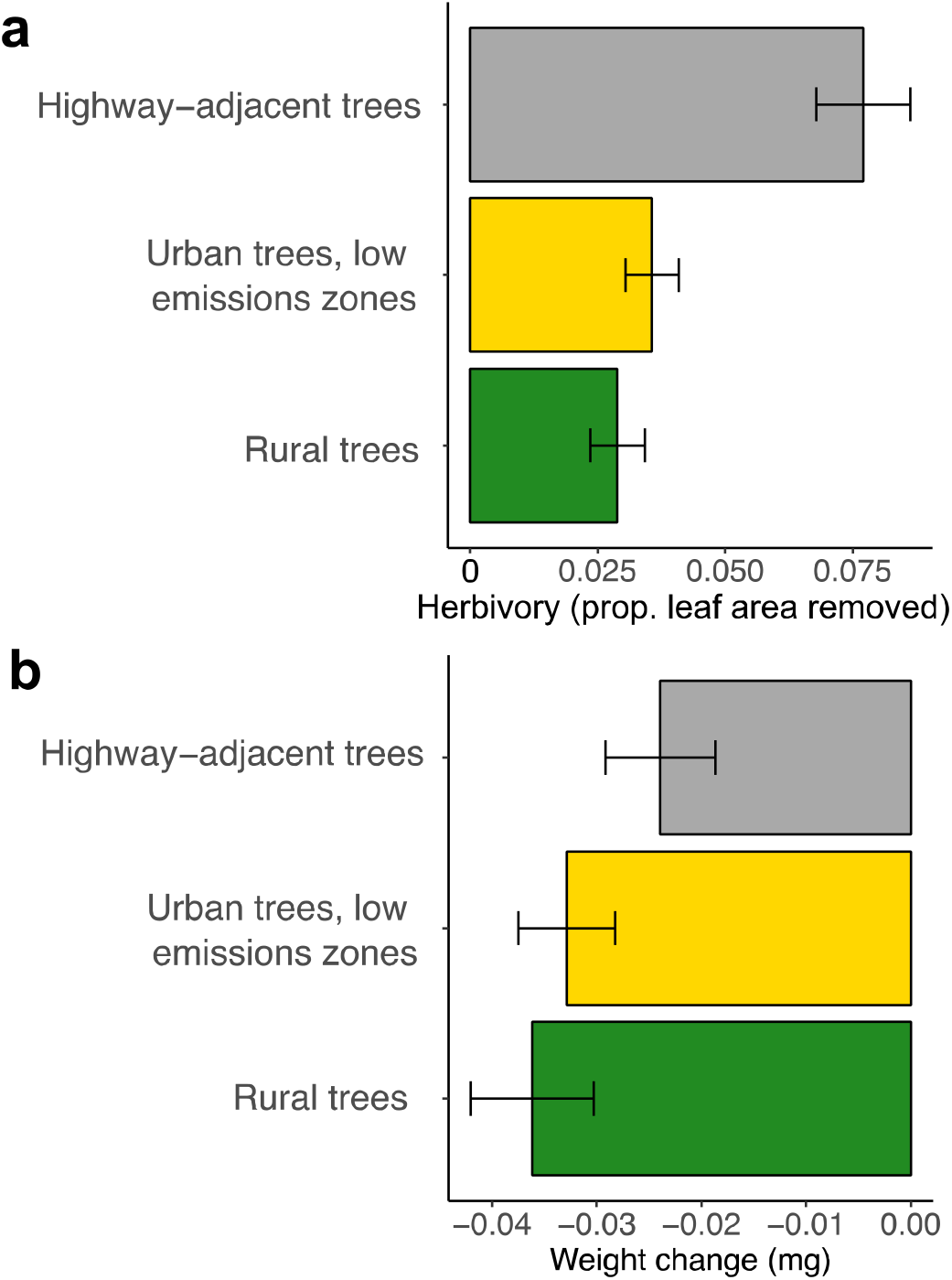
Effects of tree exposure to highways on insect damage to leaves and insect performance. a) Chewing herbivory by naive insect herbivores in the laboratory as it relates to categories of pollution exposure in trials that gave insects the choice to feed on leaves within the category of their choosing, and b) weight change in caterpillars in no-choice trials fed leaves from one pollution exposure category.

## Discussion

Vehicle pollution was strongly associated with elevated insect chewing herbivory across all models. Trees in high-pollution areas displayed more chewing herbivory than trees in urban areas with low levels of pollution or rural areas: On highway-adjacent trees, leaves from branches bordering highways were eaten more than branches not bordering highways. In laboratory assays, leaves from highway-adjacent trees were both more attractive and more nutritious for generalist caterpillars, providing evidence for plant-mediated elevation of insect chewing herbivory in response to on-road pollution. Effects of pollution on leaf mining, in contrast, were weaker but trended toward less leaf mining in more polluted areas.

We assessed effects of pollution on herbivory using four metrics: a categorical representation of how focal trees were selected (from rural, highway-adjacent, urban low emissions, and urban high emissions sites), a continuous metric of tree proximity to highways, a categorical metric of whether a branch was collected from a side of a tree bordering a highway, and—the metric that arguably serves as the most direct measure of pollution from vehicles—on-road emissions from the DARTE database (Gately et al., 2015). Across every model assessing effects of these predictors, pollution was associated with elevated leaf chewing, and differences between these predictors reveal additional information about the specific drivers of herbivory. On-road pollution derived from the DARTE database had the strongest effect of any continuous predictor, supporting our hypothesis that vehicle pollution is a key driver of insect chewing on leaves. Effects of distance from a major road on herbivory were slightly weaker and more variable, but trees nearer to highways still clearly displayed more herbivory than those further away, suggesting that distance from highways may serve as an adequate proxy for tree exposure to on-road pollution. This proxy could be useful in a management context—for instance, if on-road emissions data are not available to aid in decision making about where to plant herbivory-resistant tree species and cultivars.

The metric that explained the most variation in chewing herbivory was sampling category, such that highway-adjacent trees displayed more herbivory than any other trees, and rural trees experienced the lowest amount of herbivory. Urban trees from high-and low-emissions zones displayed herbivory levels that fell between these other two categories, though ‘high emissions’ trees did not display higher herbivory than ‘low emissions’ trees. This might be because, in some cases, trees may not fall into these categories reliably on-the-ground because DARTE data are at a one-by-one-km resolution, and levels of pollution can vary more locally. Elevated levels of on-road pollutants are often localized to a few hundred meters from roadways, at most, because pollutants are quickly dispersed (Brugge et al., 2007, but see Cobley & Pataki, 2019). Highly localized pollution is consistent with the strong effects of highway-adjacency on herbivory that we documented. We note, however, that even though DARTE may not capture on-road pollution reliably when included categorically, it was representative enough in our models to serve as a strong continuous predictor of herbivory.

Low amounts of herbivory at rural trees may indicate that both pollution and other factors drive differences in herbivory differences between urban and rural sites. Our models accounted for NDVI as a measure of surrounding vegetation, and chewing herbivory on trees from rural areas was still statistically distinguishable from the three urban sampling categories; thus, we suggest that rural sites have additional factors protecting trees from high levels of mandibulate herbivory, such as elevated biological control in rural compared to urban habitats (Raupp et al., 2010; Shrewsbury & Raupp, 2006) and/or above or belowground conditions that aid in tree defense against herbivores (Moreira et al., 2019).

Along these lines, laboratory assays demonstrated that rural trees are less palatable and are of poorer nutritional quality for caterpillars than leaves from urban, highway-adjacent habitats. Without these assays, it would be impossible to tell whether higher herbivory rates on polluted trees were a result of higher or lower leaf quality at those sites; more feeding on polluted plants could indicate that leaves were more nutritious for herbivores, or less, i.e., that individual insects had to eat more polluted leaf material to meet their nutritional needs (Lincoln et al., 1986, 1993; Schäedler et al., 2007). Contrary to this possibility, lower weight loss by highway-adjacent caterpillars in laboratory assays indicates that leaves from polluted sites are actually of higher quality for herbivores. Though caterpillars were attracted to leaves from highway-adjacent trees and lost less weight when fed those leaves, all caterpillars in the laboratory trial lost weight, which is not surprising; *Q. lobata* is a novel food source for this caterpillar population, and its leaves are much tougher than those of their host lupine. Our focal caterpillar species exhibits very limited dispersal and gene flow, and it might as a result be locally adapted to its host plant even though it can feed on other species (Harrison, 1997).

Together, our data support a scenario in which leaves from highway adjacent trees are both more attractive and nutritious to generalist herbivores, and fitness of these herbivores may be higher on polluted trees. However, more herbivory on more polluted trees could result from multiple proximate mechanisms between which we cannot distinguish in this study; higher herbivory on highway-adjacent trees, for instance, may be a result of unique insect communities at these sites, higher feeding rates by individual insects, higher abundance due to the attractiveness of leaves to adult moths who lay eggs, and/or higher abundance due to elevated insect fitness on more polluted trees. Our study provides evidence for the potential importance of the latter three mechanisms.

We did not find evidence that on-road pollution is associated with changes in elemental nutrients that could explain patterns in herbivory. Instead, it is possible that pollution affects specific nutritional or defensive compounds, and future studies will be required to identify these compounds. It is also possible that elemental leaf nutrient changes at different points during the season are affected by vehicle pollution, but that these changes were masked by effects of the herbivores on elemental leaf composition. For instance, leaf miners may artificially elevate leaf nitrogen content. We chose minimally damaged leaves for nutrient analyses, and we therefore find this explanation unlikely. Instead, we suspect that vehicle pollution depresses defensive pathways within trees and reduces the concentrations of key compounds that protect against herbivore damage (Moreira et al., 2019). Evidence for this scenario emerged in laboratory assays, wherein caterpillars only very rarely chose to feed on leaves from rural trees in ‘choice’ trials. Moreover, even when rural leaves were the only food available to caterpillars in ‘no choice’ trials, they avoided feeding on rural leaves, a mechanism that might have driven more weight loss in these caterpillars compared to those fed leaves from highway-adjacent trees.

We included a wide range of measured covariates in models to test alternative hypotheses about what may drive patterns of herbivory on urban trees across pollution gradients. Perhaps surprisingly, light pollution, surface temperature, and tree water stress were not implicated as drivers of herbivory on urban *Q. lobata*. Accounting for these variables provided more support that on-road pollution is a key mechanism driving herbivory. Tree size and NDVI were the only covariates that explained additional variation in chewing herbivory, such that bigger trees and trees surrounded by less vegetation were eaten more by chewing herbivores (though no covariates affected leaf mining). Our work adds to mounting evidence that surrounding vegetation has a protective effect on trees, potentially through the provision of biological control services that are supported by additional plant cover, diversity, and/or complexity (Korányi et al., 2022; Nighswander et al., 2021; Philpott & Bichier, 2017; Shrewsbury & Raupp, 2006). We suspect that larger trees were eaten more than smaller trees because they are older and have therefore had more years to accumulate herbivores that are sedentary and tend to mate and lay eggs on the same tree year after year. Larger trees may also be more apparent to herbivores (Feeny, 1976).

Though our study points to the key importance of bottom-up processes on insect herbivory across urban pollution gradients, top-down effects may also partially explain why highway-adjacent leaves were eaten more. Airborne pollutants may disrupt or dilute chemical signals to predators and parasitoids (Blande, 2021). Highway-associated noise could also discourage foraging by birds, potentially because traffic noise precludes their communication or otherwise deters them from being present on highway-adjacent trees (Grade & Sieving, 2016). More studies are also needed to determine if top-down processes work in tandem with bottom-up processes we identified to elevate chewing herbivory on polluted trees.

Chewing herbivory was lower in 2021 than in 2020 despite lower urban vehicle traffic across the USA in 2020 that has been linked to reduced pollution (Pitiranggon et al., 2022). This pattern that at first glance is contrary to our hypothesis that chewing herbivory is elevated on some trees compared to others because of on-road pollution. While differences in pollution exposure among trees located in different areas may predict levels of herbivory, other factors may be more important in driving how much plants are eaten from year to year. This underscores the need for more long-term studies on insect herbivory in cities to promote a temporally explicit understanding of the major drivers of plant-insect herbivore interactions (Ossola et al., 2021).

Regardless, our study highlights the importance of planting decisions along major roadways. The concept of “right tree, right place” has long stated that tree selection should be aimed at maximizing the performance in urban areas (Minckler, 1941; Morakinyo et al., 2020; Wang et al., 2022). *Quercus lobata* and other species that are highly susceptible to herbivores may provide ecosystem services sub-optimally along highways, and may have shorter lifespans due to chronic damage promoted by on-road pollution (Pearse et al., 2015). Identifying tree species that are robust to pollution, and resistant to insects that may benefit from pollution, could be a novel consideration in planting decisions. This consideration may become even more important as many cities become drier and hotter, and insect herbivores have disproportionate impacts on tree growth (Meineke & Frank, 2018). Because city-owned trees are planted and cannot themselves evolve in response to climate change, we may be required to develop new cultivars to promote robust trees along roadways.

Our research also has implications for other cultivated and wild plant communities worldwide. In most countries, it is difficult to locate land that is more than one mile from a roadway (Riitters & Wickham, 2003). Initiatives such as those spearheaded by the former First Lady Claudia “Lady Bird” Johnson to plant native wildflowers along state and federal highways (Gould, 1996) have the potential to contribute to local biodiversity (O’Sullivan et al., 2017). However, these same plants may take up pollutants from vehicles, with unknown consequences for interactions between plants, insects, and higher trophic levels. Our research suggests that on-road pollution ca have consequences for plant-insect interactions, which deserve to be studied further in the contexts of plant performance and services, biological conservation, and climate change. Our research also adds to a now growing chorus of studies (Lahr et al., 2018; Youngsteadt et al., 2015, 2017) demonstrating the scientific value of intra-urban gradients of particular variables (heat, pollution, surrounding vegetation). These specific gradients move beyond urban-to-rural gradients to isolate particular mechanisms shaping ecological functions in rural and urban habitats.

## Authors’ contributions

EKM conceived of the urban field study, collaborated with RK and DSE on design of the palatability studies, led statistical analyses, and co-wrote the manuscript. DSE collected data, helped manage undergraduate research activities, and helped edit the manuscript. RK helped design and collect data for palatability studies and co-wrote the manuscript.

## Data availability statement

Upon publication, all data from this project will be made available on the Dryad database with a corresponding DOI. All code and data will additionally be made available on the Meineke Lab website at a designated URL.

## Acknowledgements

We thank the City of Sacramento for providing a shapefile of city-owned urban trees and for providing permissions and permits that made this research possible. This study was funded by the UC Davis Department of Entomology and Nematology.

