## Appendices for "Vehicle pollution is associated with elevated insect damage to street trees"

**Supplementary Figures, Figure Captions, and Tables**

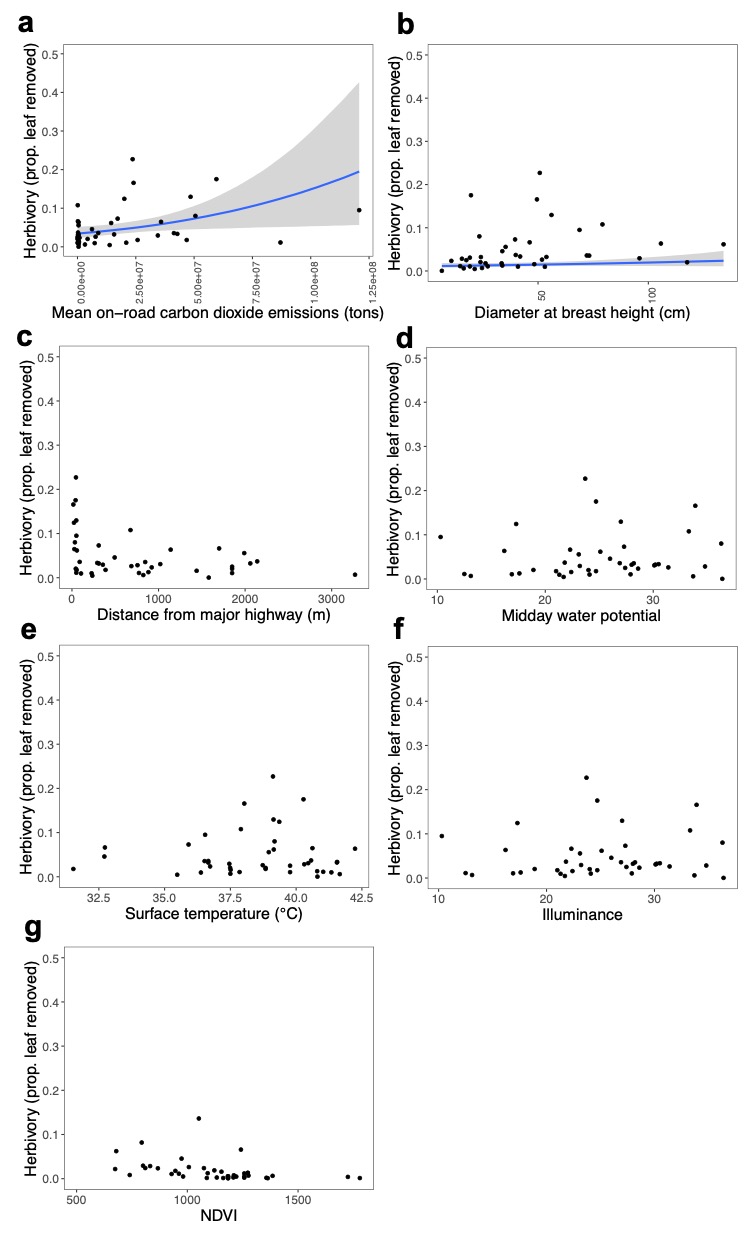

**Fig. S1. Investigation of various potential mechanisms driving insect chewing herbivory on focal trees.** Data shown are from 2020, and plots from 2021 appear similar. Model fits are only shown for posterior distributions with 95% credible intervals not crossing zero.

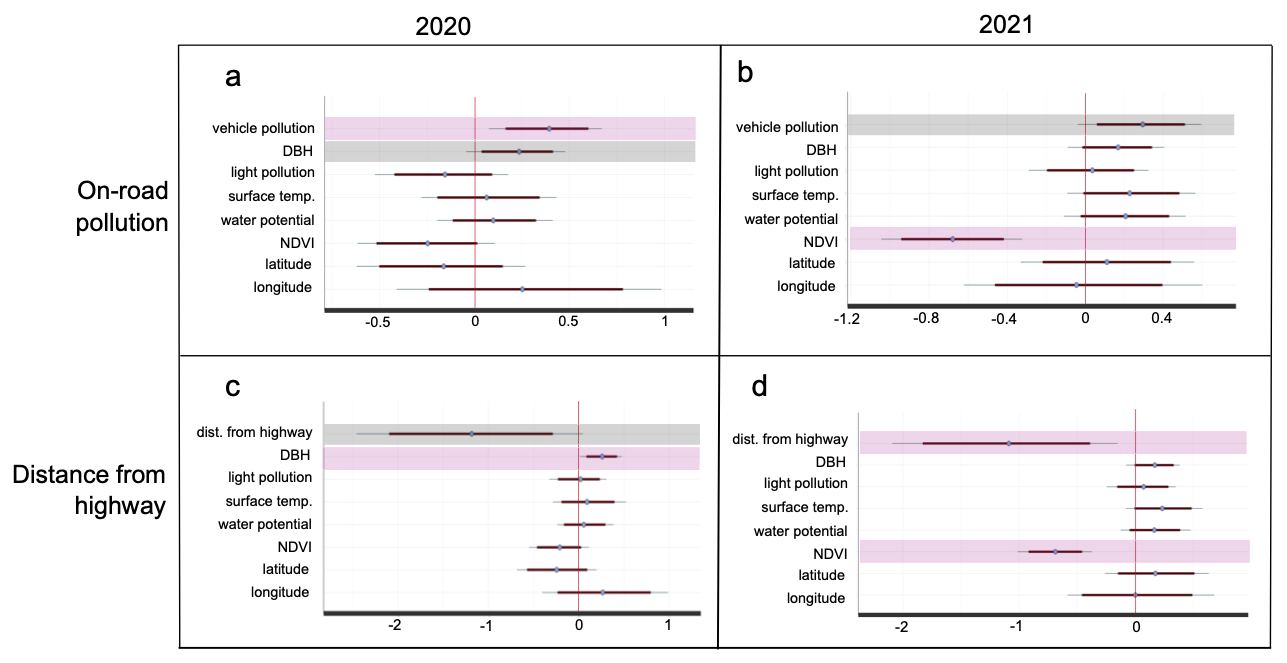

**Fig. S2.** Model estimates showing the predicted effects of pollution, location, and environmental variables on insect chewing herbivory by year. Bold lines represent 85% credibility intervals, and narrow lines represent 95% credibility intervals. Predictors with 85% credibility intervals not crossing zero are highlighted in grey, and those with 95% credibility intervals not crossing zero are highlighted in purple.

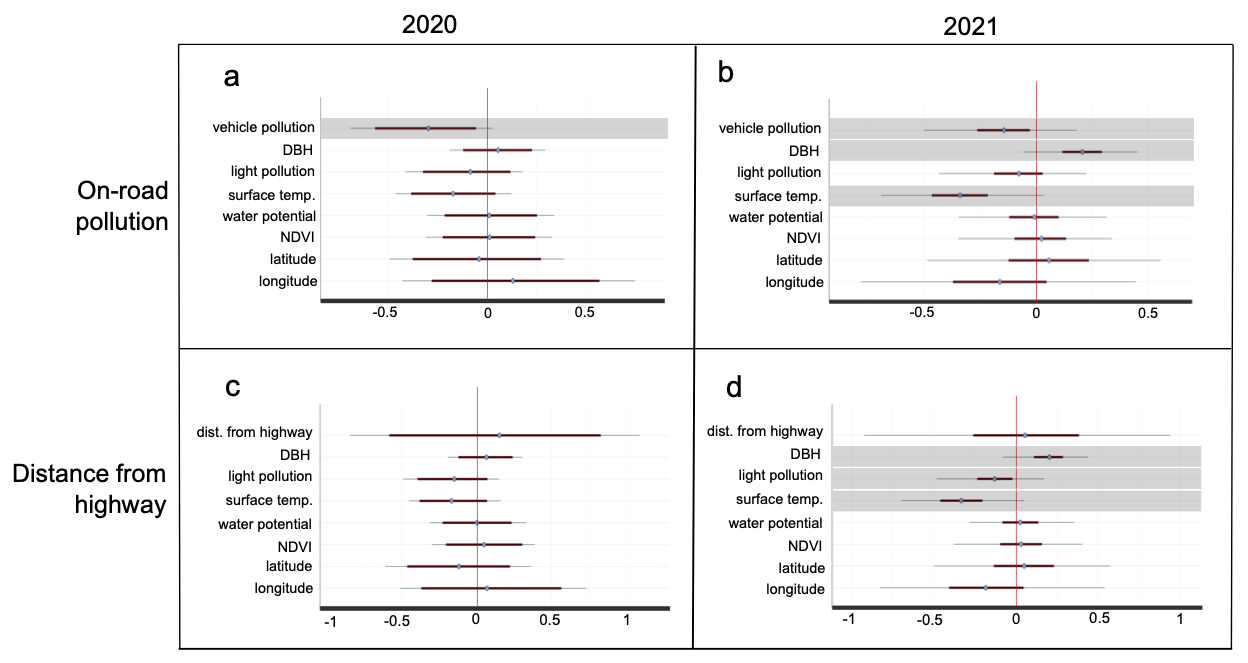

**Fig. S3.** Model estimates showing the predicted effects of pollution, location, and environmental variables on insect mining herbivory. Bold lines represent 85% credibility intervals, and narrow lines represent 95% credibility intervals. Predictors with 85% credibility intervals not crossing zero are highlighted in grey.

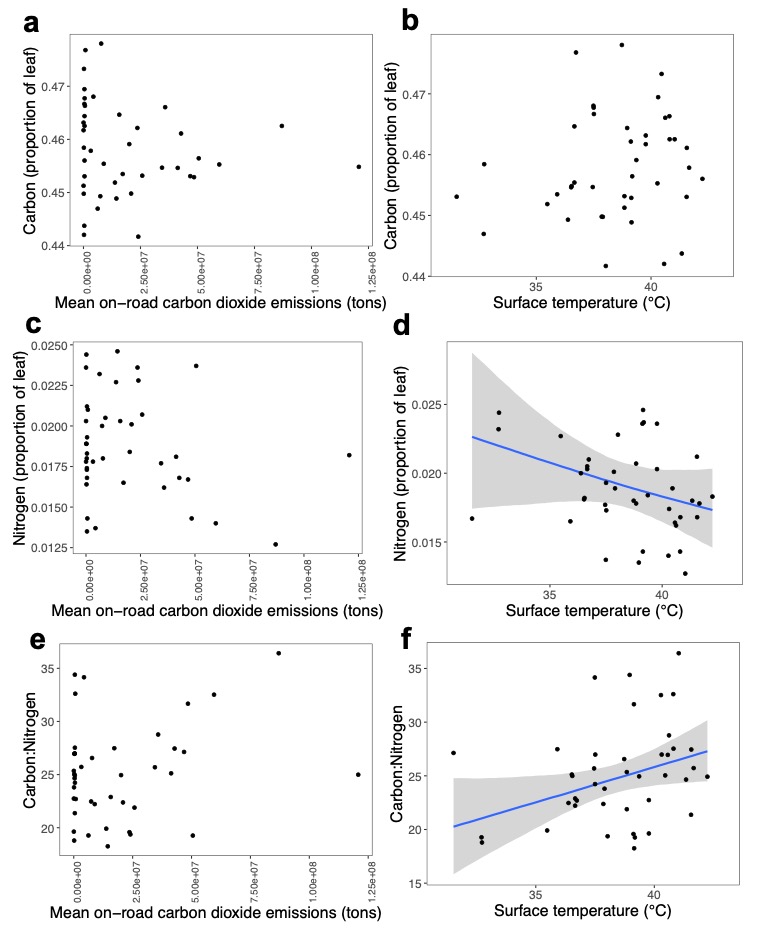

**Fig. S4.** Carbon, nitrogen, and carbon:nitrogen concentrations as they relate to pollution and surface temperature. Fitted lines are only included for predictors with posterior distributions not crossing zero or only slightly crossing zero (in the case of d, whose statistics are included in Table S4).

**Table S1. Effects of key predictors on insect chewing herbivory in 2020.** Bayesian models predicting the effects of a) vehicle emissions category, b) distance from the nearest major highway, c) on-road CO_2_ emissions, along with covariates included in all models: light pollution (illuminance), surface temperature, stem water potential (tree water stress), tree size (diameter at breast height), and latitude and longitude of individual focal trees on proportion of leaf area removed by mandibulate herbivores during the 2020 growing season. For each parameter, *ß*_avg_ is the estimated average effect on insect chewing herbivory. Values of each continuous variable were scaled prior to analysis. Thus, larger deviations of *ß*_avg_ values from zero indicate a larger effect of the parameter, and these effects can be compared across parameters. The effective sample size is indicated by n_eff_, and Rhat = 1 indicates convergence.

***a. Model with pollution exposure categories (R^2^= 0.55)***

| **Variable** | *ß*_avg_ | **SE** | **2.5%** | **97.5%** | **n_eff_** | **Rhat** |
| --- | --- | --- | --- | --- | --- | --- |
| Intercept | -3.54 | 0.56 | -4.68 | -2.46 | 1781 | 1.00 |
| Highway-adjacent | 1.61 | 0.32 | 0.96 | 2.24 | 2431 | 1.00 |
| Urban, low emissions | 0.65 | 0.35 | -0.05 | 1.33 | 2207 | 1.00 |
| Rural | -0.43 | 1.24 | -2.81 | 2.07 | 1787 | 1.00 |
| Illuminance | -0.07 | 0.17 | -0.42 | 0.26 | 2883 | 1.00 |
| Surface temperature | -0.20 | 0.17 | -0.53 | 0.14 | 2508 | 1.00 |
| Water potential | 0.15 | 0.14 | -0.13 | 0.42 | 2568 | 1.00 |
| Tree size | 0.19 | 0.11 | -0.03 | 0.48 | 2388 | 1.00 |
| Latitude | -0.05 | 0.26 | -0.53 | 0.40 | 2291 | 1.00 |
| Longitude | -0.32 | 1.09 | -2.46 | 1.86 | 1663 | 1.00 |
| NDVI | -0.32 | 0.15 | -0.62 | -0.02 | 2883 | 1.00 |

***b. Mechanistic model including on-road vehicle emissions (R^2^= 0.27)***

| **Variable** | *ß*_avg_ | **SE** | **2.5%** | **97.5%** | **n_eff_** | **Rhat** |
| --- | --- | --- | --- | --- | --- | --- |
| Intercept | -3.14 | 0.23 | -3.60 | -2.70 | 2769 | 1.00 |
| On-road CO_2_ emissions | 0.39 | 0.16 | 0.08 | 0.67 | 2553 | 1.00 |
| Illuminance | -0.16 | 0.18 | -0.53 | 0.19 | 2798 | 1.00 |
| Surface temperature | 0.06 | 0.18 | -0.28 | 0.43 | 3139 | 1.00 |
| Water potential | 0.10 | 0.15 | -0.19 | 0.40 | 3148 | 1.00 |
| Tree size | 0.23 | 0.13 | -0.03 | 0.48 | 3064 | 1.00 |
| Latitude | -0.17 | 0.22 | -0.61 | 0.25 | 3334 | 1.00 |
| Longitude | 0.27 | 0.34 | -0.37 | 0.98 | 2770 | 1.00 |
| NDVI | -0.24 | 0.19 | -0.61 | 0.12 | 3009 | 1.00 |

***c. Mechanistic model including distance from nearest major highway (R^2^= 0.26)***

| **Variable** | *ß*_avg_ | **SE** | **2.5%** | **97.5%** | **n_eff_** | **Rhat** |
| --- | --- | --- | --- | --- | --- | --- |
| Intercept | -3.59 | 0.35 | -4.29 | -2.89 | 3139 | 1.00 |
| Distance from highway | -1.20 | 0.64 | -2.48 | 0.14 | 3071 | 1.00 |
| Illuminance | 0.01 | 0.15 | -0.30 | 0.29 | 3221 | 1.00 |
| Surface temperature | 0.09 | 0.20 | -0.29 | 0.50 | 3479 | 1.00 |
| Water potential | 0.06 | 0.16 | -0.24 | 0.39 | 3438 | 1.00 |
| Tree size | 0.25 | 0.12 | 0.01 | 0.48 | 4012 | 1.00 |
| Latitude | -0.24 | 0.23 | -0.68 | 0.20 | 2982 | 1.00 |
| Longitude | 0.28 | 0.36 | -0.39 | 1.00 | 2831 | 1.00 |
| NDVI | -0.21 | 0.17 | -0.53 | 0.11 | 3274 | 1.00 |

**Table S2. Effects of key predictors on insect chewing herbivory in 2021.** Bayesian models predicting the effects of a) vehicle emissions category, b) distance from the nearest major highway, c) on-road CO_2_ emissions, along with covariates included in all models: light pollution (illuminance), surface temperature, stem water potential (tree water stress), tree size (diameter at breast height), and latitude and longitude of individual focal trees on proportion of leaf area taken up by leaf mines. For each parameter, *ß*_avg_ is the estimated average effect on insect leaf mining. Values of each continuous variable were scaled prior to analysis. Thus, larger deviations of *ß*_avg_ values from zero indicate a larger effect of the parameter, and these effects can be compared across parameters. The effective sample size is indicated by n_eff_, and Rhat = 1 indicates convergence.

***a. Model with pollution exposure categories (R^2^= 0.42)***

| **Variable** | *ß*_avg_ | **SE** | **2.5%** | **97.5%** | **n_eff_** | **Rhat** |
| --- | --- | --- | --- | --- | --- | --- |
| Intercept | -4.26 | 0.35 | -4.95 | -3.60 | 2134 | 1.00 |
| Highway-adjacent | 1.11 | 0.32 | 0.45 | 1.71 | 2264 | 1.00 |
| Urban, low emissions | -0.07 | 0.37 | -0.80 | 0.68 | 2582 | 1.00 |
| Rural | -0.27 | 0.50 | -1.34 | 0.67 | 1969 | 1.00 |
| Illuminance | 0.11 | 0.16 | -0.21 | 0.41 | 2443 | 1.00 |
| Surface temperature | 0.13 | 0.19 | -0.22 | 0.52 | 2226 | 1.00 |
| Water potential | 0.26 | 0.14 | -0.05 | 0.54 | 2906 | 1.00 |
| Tree size | 0.22 | 0.12 | -0.01 | 0.43 | 2756 | 1.00 |
| Latitude | -0.02 | 0.31 | -0.62 | 0.61 | 1765 | 1.00 |
| Longitude | -0.19 | 0.45 | -1.15 | 0.60 | 2240 | 1.00 |
| NDVI | -0.81 | 0.16 | -0.12 | -0.50 | 2661 | 1.00 |

***b. Mechanistic model including on-road vehicle emissions (R^2^= 0.26)***

| **Variable** | *ß*_avg_ | **SE** | **2.5%** | **97.5%** | **n_eff_** | **Rhat** |
| --- | --- | --- | --- | --- | --- | --- |
| Intercept | -3.97 | 0.20 | -4.38 | -3.57 | 2859 | 1.00 |
| On-road CO_2_ emissions | 0.29 | 0.16 | -0.02 | 0.59 | 3001 | 1.00 |
| Illuminance | 0.03 | 0.16 | -0.29 | 0.33 | 3400 | 1.00 |
| Surface temperature | 0.24 | 0.17 | -0.08 | 0.59 | 3003 | 1.00 |
| Water potential | 0.20 | 0.16 | -0.12 | 0.51 | 3348 | 1.00 |
| Tree size | 0.17 | 0.13 | -0.09 | 0.41 | 3589 | 1.00 |
| Latitude | 0.11 | 0.23 | -0.33 | 0.56 | 3286 | 1.00 |
| Longitude | -0.04 | 0.31 | -0.63 | 0.59 | 3080 | 1.00 |
| NDVI | -0.68 | 0.18 | -1.05 | -0.33 | 2584 | 1.00 |

***c. Mechanistic model including distance from nearest major highway (R^2^= 0.28)***

| **Variable** | *ß*_avg_ | **SE** | **2.5%** | **97.5%** | **n_eff_** | **Rhat** |
| --- | --- | --- | --- | --- | --- | --- |
| Intercept | -4.34 | 0.28 | -4.93 | -3.82 | 2760 | 1.00 |
| Distance from highway | -1.12 | 0.49 | -2.12 | -0.17 | 3218 | 1.00 |
| Illuminance | 0.06 | 0.15 | -0.25 | 0.34 | 2711 | 1.00 |
| Surface temperature | 0.24 | 0.17 | -0.09 | 0.57 | 2907 | 1.00 |
| Water potential | 0.17 | 0.15 | -0.12 | 0.45 | 3113 | 1.00 |
| Tree size | 0.16 | 0.12 | -0.07 | 0.38 | 3716 | 1.00 |
| Latitude | 0.19 | 0.22 | -0.26 | 0.61 | 3267 | 1.00 |
| Longitude | 0.02 | 0.33 | -0.60 | 0.69 | 3136 | 1.00 |
| NDVI | -0.69 | 0.16 | -1.01 | -0.37 | 2801 | 1.00 |

**Table S3. Effects of key predictors on insect leaf mining herbivory in 2020.** Bayesian models predicting the effects of a) vehicle emissions category, b) distance from the nearest major highway, c) on-road CO_2_ emissions, along with covariates included in all models: light pollution (illuminance), surface temperature, stem water potential (tree water stress), tree size (diameter at breast height), and latitude and longitude of individual focal trees on proportion of leaf area taken up by leaf mines. For each parameter, *ß*_avg_ is the estimated average effect on insect leaf mining. Values of each continuous variable were scaled prior to analysis. Thus, larger deviations of *ß*_avg_ values from zero indicate a larger effect of the parameter, and these effects can be compared across parameters. The effective sample size is indicated by n_eff_, and Rhat = 1 indicates convergence.

***a. Model with pollution exposure categories (R^2^= 0.22)***

| **Variable** | *ß*_avg_ | **SE** | **2.5%** | **97.5%** | **n_eff_** | **Rhat** |
| --- | --- | --- | --- | --- | --- | --- |
| Intercept | -2.72 | 0.58 | -3.90 | -1.61 | 1817 | 1.00 |
| Highway-adjacent | -0.63 | 0.39 | -1.39 | 0.13 | 2758 | 1.00 |
| Urban, low emissions | 0.31 | 0.32 | -0.39 | 0.95 | 2661 | 1.00 |
| Rural | -0.99 | 1.21 | -3.31 | 1.44 | 1800 | 1.00 |
| Illuminance | -0.04 | 0.16 | -0.36 | 0.26 | 2337 | 1.00 |
| Surface temperature | -0.15 | 0.15 | -0.43 | 0.13 | 2815 | 1.00 |
| Water potential | 0.12 | 0.17 | -0.22 | 0.45 | 2653 | 1.00 |
| Tree size | 0.06 | 0.12 | -0.18 | 0.29 | 3769 | 1.00 |
| Latitude | 0.06 | 0.23 | -0.40 | 0.53 | 2543 | 1.00 |
| Longitude | -0.86 | 1.04 | -2.88 | 1.15 | 1732 | 1.00 |
| NDVI | 0.07 | 0.17 | -0.26 | 0.40 | 2833 | 1.00 |

***b. Mechanistic model including on-road vehicle emissions (R^2^= 0.18)***

| **Variable** | *ß*_avg_ | **SE** | **2.5%** | **97.5%** | **n_eff_** | **Rhat** |
| --- | --- | --- | --- | --- | --- | --- |
| Intercept | -3.21 | 0.18 | -3.56 | -2.86 | 3315 | 1.00 |
| On-road CO_2_ emissions | -0.31 | 0.18 | -0.68 | 0.02 | 3578 | 1.00 |
| Illuminance | -0.08 | 0.15 | -0.40 | 0.20 | 3159 | 1.00 |
| Surface temperature | -0.18 | 0.14 | -0.45 | 0.11 | 3731 | 1.00 |
| Water potential | 0.02 | 0.16 | -0.29 | 0.34 | 3231 | 1.00 |
| Tree size | 0.06 | 0.12 | -0.17 | 0.28 | 4267 | 1.00 |
| Latitude | -0.05 | 0.22 | -0.49 | 0.38 | 3336 | 1.00 |
| Longitude | 0.13 | 0.30 | -0.43 | 0.75 | 3452 | 1.00 |
| NDVI | 0.01 | 0.17 | -0.31 | 0.35 | 3326 | 1.00 |

***c. Mechanistic model including distance from nearest major highway (R^2^= 0.14)***

| **Variable** | *ß*_avg_ | **SE** | **2.5%** | **97.5%** | **n_eff_** | **Rhat** |
| --- | --- | --- | --- | --- | --- | --- |
| Intercept | -3.14 | 0.26 | -3.67 | -2.67 | 3518 | 1.00 |
| Distance from highway | 0.15 | 0.49 | -0.83 | 1.13 | 3651 | 1.00 |
| Illuminance | -0.15 | 0.16 | -0.48 | 0.15 | 3344 | 1.00 |
| Surface temperature | -0.15 | 0.15 | -0.44 | 0.15 | 3909 | 1.00 |
| Water potential | 0.01 | 0.16 | -0.30 | 0.33 | 3492 | 1.00 |
| Tree size | 0.07 | 0.13 | -0.19 | 0.31 | 4004 | 1.00 |
| Latitude | -0.12 | 0.24 | -0.60 | 0.34 | 3603 | 1.00 |
| Longitude | 0.09 | 0.32 | -0.52 | 0.75 | 3045 | 1.00 |
| NDVI | 0.10 | 0.33 | -0.50 | 0.76 | 3945 | 1.00 |

**Table S4. Effects of key predictors on insect leaf mining herbivory in 2021.** Bayesian models predicting the effects of a) vehicle emissions category, b) distance from the nearest major highway, c) on-road CO_2_ emissions, along with covariates included in all models: light pollution (illuminance), surface temperature, stem water potential (tree water stress), tree size (diameter at breast height), and latitude and longitude of individual focal trees on proportion of leaf area taken up by leaf mines. For each parameter, *ß*_avg_ is the estimated average effect on insect leaf mining. Values of each continuous variable were scaled prior to analysis. Thus, larger deviations of *ß*_avg_ values from zero indicate a larger effect of the parameter, and these effects can be compared across parameters. The effective sample size is indicated by n_eff_, and Rhat = 1 indicates convergence.

***a. Model with pollution exposure categories (R^2^= 0.23)***

| **Variable** | *ß*_avg_ | **SE** | **2.5%** | **97.5%** | **n_eff_** | **Rhat** |
| --- | --- | --- | --- | --- | --- | --- |
| Intercept | -5.73 | 0.39 | -6.50 | -4.96 | 2522 | 1.00 |
| Highway-adjacent | 0.19 | 0.41 | -0.67 | 0.97 | 2691 | 1.00 |
| Urban, low emissions | 0.00 | 0.39 | -0.75 | 0.74 | 2711 | 1.00 |
| Rural | -0.03 | 0.54 | -1.13 | 0.99 | 2447 | 1.00 |
| Illuminance | -0.14 | 0.18 | -0.49 | 0.19 | 3034 | 1.00 |
| Surface temperature | -0.32 | 0.22 | -0.74 | 0.14 | 2501 | 1.00 |
| Water potential | 0.04 | 0.18 | -0.32 | 0.40 | 2572 | 1.00 |
| Tree size | 0.20 | 0.13 | -0.09 | 0.44 | 3400 | 1.00 |
| Latitude | 0.01 | 0.36 | -0.67 | 0.72 | 2375 | 1.00 |
| Longitude | -0.25 | 0.48 | -1.25 | 0.62 | 2248 | 1.00 |
| NDVI | -0.01 | 0.20 | -0.40 | 0.37 | 2899 | 1.00 |

***b. Mechanistic model including on-road vehicle emissions (R^2^= 0.25)***

| **Variable** | *ß*_avg_ | **SE** | **2.5%** | **97.5%** | **n_eff_** | **Rhat** |
| --- | --- | --- | --- | --- | --- | --- |
| Intercept | -5.71 | 0.24 | -6.18 | -5.26 | 3072 | 1.00 |
| On-road CO_2_ emissions | -0.15 | 0.17 | -0.51 | 0.16 | 3305 | 1.00 |
| Illuminance | -0.08 | 0.17 | -0.43 | 0.22 | 3718 | 1.00 |
| Surface temperature | -0.34 | 0.19 | -0.69 | 0.02 | 3731 | 1.00 |
| Water potential | -0.01 | 0.17 | -0.35 | 0.32 | 4341 | 1.00 |
| Tree size | 0.21 | 0.13 | -0.06 | 0.44 | 3728 | 1.00 |
| Latitude | 0.06 | 0.27 | -0.48 | 0.58 | 4222 | 1.00 |
| Longitude | -0.16 | 0.32 | -0.78 | 0.48 | 3434 | 1.00 |
| NDVI | 0.02 | 0.18 | -0.35 | 0.35 | 3634 | 1.00 |

***c. Mechanistic model including distance from nearest major highway (R^2^= 0.29)***

| **Variable** | *ß*_avg_ | **SE** | **2.5%** | **97.5%** | **n_eff_** | **Rhat** |
| --- | --- | --- | --- | --- | --- | --- |
| Intercept | -5.70 | 0.28 | -6.27 | -5.18 | 3486 | 1.00 |
| Distance from highway | 0.05 | 0.45 | -0.84 | 0.92 | 3955 | 1.00 |
| Illuminance | -0.13 | 0.16 | -0.47 | 0.16 | 2999 | 1.00 |
| Surface temperature | -0.33 | 0.20 | -0.70 | 0.08 | 3618 | 1.00 |
| Water potential | 0.03 | 0.16 | -0.28 | 0.34 | 4079 | 1.00 |
| Tree size | 0.20 | 0.14 | -0.08 | 0.46 | 3775 | 1.00 |
| Latitude | 0.03 | 0.27 | -0.50 | 0.58 | 3872 | 1.00 |
| Longitude | -0.16 | 0.33 | -0.79 | 0.50 | 2798 | 1.00 |
| NDVI | 0.03 | 0.18 | -0.34 | 0.36 | 3443 | 1.00 |

**Table S5. Key predictors of leaf nutrient content.** Of all key predictors included in models—on-road CO_2_ emissions, light pollution (illuminance), surface temperature, stem water potential (tree water stress), tree size (diameter at breast height), and latitude and longitude of individual trees—selected statistics are listed for predictors with posterior distributions not crossing zero or only slightly crossing zero, i.e., predictors for which evidence indicates effects on leaf nutrient content. For each parameter, *ß*_avg_ is the estimated average effect on leaf nutrients. Values of each continuous variable were scaled prior to analysis. Thus, larger deviations of *ß*_avg_ values from zero indicate a larger effect of the parameter, and these effects can be compared across parameters. The effective sample size is indicated by n_eff_, and Rhat = 1 indicates convergence.

***a. Response: Nitrogen (R^2^= 0.36)***

| **Variable** | *ß*_avg_ | **SE** | **2.5%** | **97.5%** | **n_eff_** | **Rhat** |
| --- | --- | --- | --- | --- | --- | --- |
| Surface temperature | -0.08 | 0.06 | -0.20 | 0.03 | 3839 | 1.00 |

***b. Response: Carbon:Nitrogen (R^2^= 0.33)***

| **Variable** | *ß*_avg_ | **SE** | **2.5%** | **97.5%** | **n_eff_** | **Rhat** |
| --- | --- | --- | --- | --- | --- | --- |
| Surface temperature | 2.07 | 1.00 | 0.03 | 4.07 | 3309 | 1.00 |
